# Small molecule docking of DNA repair proteins associated with cancer survival following PCNA metagene adjustment: A potential novel class of repair inhibitors

**DOI:** 10.1101/485375

**Authors:** Leif E. Peterson

## Abstract

Natural and synthetic small molecules from the NCI Developmental Therapeutics Program (DTP) were employed in molecular dynamics-based docking with DNA repair proteins whose RNA-Seq based expression was associated with overall cancer survival (OS) after adjustment for the PCNA metagene. The compounds employed were required to elicit a sensitive response (vs. resistance) in more than half of the cell lines tested for each cancer. Methodological approaches included peptide sequence alignments and homology modeling for 3D protein structure determination, ligand preparation, docking, toxicity and ADME prediction. Docking was performed for unique lists of DNA repair proteins which predict OS for AML, cancers of the breast, lung, colon, and ovaries, GBM, melanoma, and renal papillary cancer. Results indicate hundreds of drug-like and lead-like ligands with best-pose binding energies less than −6 kcal/mol. Ligand solubility for the top 20 drug-like hits approached lower bounds, while lipophilicity was acceptable. Most ligands were also blood-brain barrier permeable with high intestinal absorption rates. While the majority of ligands lacked positive prediction for Herg channel blockage and Ames carcinogenicity, there was considerable variation for predicted fathead minnow, honey bee, and *Tetrahymena pyriformis* toxicity. The computational results suggest the potential for new targets and mechanisms of repair inhibition and can be directly employed for in vitro and in vivo confirmatory laboratory experiments to identify new targets of therapy for cancer survival.

## INTRODUCTION

An important hallmark of the evolution of cancer involves upregulation of DNA repair proteins to counter the increased mutational load as a result of widespread genomic instability. Genomic instability arises from chromosomal aberrations, the accumulation of somatic mutations in driver genes, microsatellite instability, and erroneous base inclusions by polymerases, etc. [1–3]. Despite the genetic selective pressure that ensues, a tumor cell cannot survive excessive DNA damage beyond a certain mutational threshold. To exploit this phenomenon, DNA repair inhibitors have been introduced to augment molecular-based therapies for oncogene addiction and synthetic lethality [4–6]. DNA repair inhibitors for oncotherapy fall into several classes, which include poly (ADP) ribose polymerase (PARP) inhibitors, nucleotide excision repair (NER) inhibitors, DNA-PK inhibitors, MRN, ATM, ATR, CHK1/2 inhibitors, RAD51 inhibitors, and base excision repair (BER) inhibitors. PARP inhibitors have demonstrated great promise in the treatment of patients with deficiencies in homologous recombination (HR) DNA repair, such as those with loss of BRCA1 or BRCA2 function[5, 7–12]. Nucleotide excision repair inhibitors target more than thirty protein-protein interactions and removes DNA adducts caused by platinum-based chemotherapy[13–17]. DNA-PK inhibitors [18–22] target DNA-dependent protein kinase (DNA-PK) enzymes, which play a role in detection and repair of DSB via the non-homologous end-joining pathway. MRN, ATM, ATR, CHEK1/2 inhibitors [23, 24] target the kinases MRN, ATM, ATR, CHK1, and CHK2, and RAD51 [25–28] inhibitors target RAD51, a key protein of homologous recombination to repair DSB and inter-strand cross-links. BER inhibitors [29, 30] target BER proteins, which can protect a cell after endogenous or exogenous genotoxic stress, since a deficiency in BER can result in stress-induced apoptosis, necrotic cell death, mutagenesis, and chromosomal rearrangements.

In a recent investigation of TCGA RNA-Seq data and DNA repair genes, we identified sets of DNA repair genes for various cancers (Table 1) whose down-regulated expression patterns were associated with prolonged overall survival (OS)[31]. Prior to gene identification, DNA repair gene expression was adjusted for age at diagnosis, stage, and expression of the PCNA metagene[32]. Statistical randomization tests were also employed in which sets of DNA repair genes were randomly sampled for generating empirical, distribution-free, p-values. Using the list of DNA repair genes whose down-regulation was associated with longer OS, we hypothesized that compounds which strongly bind to these repair proteins could potentially establish new leads for novel DNA repair inhibitors. Additional insight could be gained from our use of the PCNA metagene to adjust expression of DNA repair genes prior to survival prediction, since this has heretofore eluded systematic investigation. Therefore, it warrants noting that the DNA repair genes in Table 1 would not have been identified without PCNA adjustment, and it is for this reason we believe this new perspective could very well define new targets for cancer therapy.

**Table 1.** DNA Repair proteins employed in ligand-receptor docking[31].

| **Cancer** | **DNA repair proteins whose downregulation is associated with prolonged OS** |
| --- | --- |
| AML | *RAD23A, EME2, APEX2* |
| Breast | *ATRIP, FANCC, RAD1, RFC3, NEIL3, EXO1, FANCB, FANCD2, FANCI, RAD51, XRCC4* |
| Colon | *RAD23A, RFC2, POLL, MLH3, FANCL* |
| GBM | *XRCC5, NBN, DDB1, GTF2H2, ERCC4, ALKBH2, APEX1, PRKDC, PMS1, REV1* |
| Renal papillary | *BLM, RAD1, FEN1, LIG1, EXO1, MSH6, BRCA2, EME1, FANCB, LIG4* |
| Lung | *BRCA1, NBN, RAD1, NEIL3, MMS19, FANCI, XRCC1, XRCC5* |
| Melanoma | *MDC1, NBN, MUTYH, POLE, UNG, FANCE, FANCI, POLI, POLK* |
| Ovarian | *PARP2, GTF2H4, SMUG1, DCLRE1B, GPS1, FANCL, APEX2, PMS1, XRCC6, MSH6, UNG, RAD51, RAD23A, EXO1, MUS81* |

The purpose of this investigation was to perform *in silico* structural drug discovery hinged to molecular dynamics (MD) to identify which compounds from the US National Cancer Institute’s (NCI) Developmental Therapeutics Program (DTP) repository tightly bind to DNA proteins identified. The DTP compounds used for ligand-receptor docking analysis will be filtered from the entire list of DTP-tested compounds, for compounds that are associated with cell line sensitivity rather than resistance. Computational toxicology and absorption, distribution, metabolism, and excretion (ADME) will also be employed to address potential safety concerns on a preliminary basis. This initial investigation can be followed by future in vitro and in vivo experiments such as qPCR and mouse/patient-derived xenografts to establish supportive lines of evidence for efficacy.

## METHODS

### Small-Molecule Ligand Library

The NCI’s DTP maintains a repository of synthetic and pure natural products that are available to investigators for non-clinical research purposes[33]. The repository collection is a uniquely diverse set of more than 200,000 compounds that have been either submitted to DTP for biological evaluation or, in some cases, synthesized under DTP auspices. Drug activity levels expressed as 50% growth inhibitory levels (GI50s) are determined by the DTP at 48 hours using the sulphorhodamine B assay[34]. All GI50 data was assessed and transformed to z-scores as described previously[35].

### Ligand Selection from DTP

Activity z-scores for 21,737 DTP compounds tested for dose-response for each cell line were obtained using the CellMiner platform (V2.1 [36]). Activity z-scores >0.5 indicate cell line sensitivity, while z-scores <0.5 imply resistance. Compound activity z-scores were available for 60 cancer cell lines representing the following cancers: breast (5 cell lines), CNS(6), colon(7), leukemia(6), melanoma(10), lung(9), ovarian(7), prostate(2), and renal(8). For each cancer, a compound was selected if the z-score >0.5 for more than half of the cell lines available.

### Protein (receptor) 3D structures

The receptors (proteins) employed in this investigation (Table 1) were identified during our previous work with TCGA RNA-Seq data and the PCNA metagene to have an association with prolonged overall survival (OS) when down-regulated. FASTA amino acid sequences (*Homo Sapiens*) for these proteins were obtained from Uniprot [37]. Homology modeling for obtaining a consensus 3D protein structure was determined using Swiss Model [38–40], which is based on Qmean [41], quaternary structure prediction/ QSQE[42], BLAST [43, 44], and the BLOSUM amino acid substitution matrix[45, 46], and were saved in PDB format. Molecular charges were merged, and non-polar hydrogens, lone pairs, and water molecules were removed using the.NET assembly of OpenBabel (OB) [47].

### Molecular Ligand-Receptor Docking

Ligand-receptor docking is an MD approach for reproducing chemical potentials which determine the bound conformation preference and free energy of binding between a ligand and its receptor. The MD technique seeks to establish the optimal receptor binding pocket (pose) with a minima in the energy profile, shape, and temperature [48], while assuming consistency in the ligand charge distribution and protonation states for the bound and unbound forms. At each receptor pocket identified, several poses are evaluated while iterating through alternative conformations of the ligand at its rotatable covalent bonds.

Prior to docking, ligand SMILES strings were converted to 3-dimensional SDF format containing partial charges of each atom. The .NET OB assembly was used for adding hydrogens to ligands and performing energy minimization of ligands and receptors using the Merck MMFF94 force field [49], with 250 iterations during conjugate gradient convergence. Energy-minimized ligands and receptors were saved in PDBQT format. Vina [50] was used for ligand-receptor docking on Amazon AWS cloud formations with Linux high-performance compute clusters. A total of 10 ligand poses were assessed at each receptor pocket identified, and the best pose was assumed to have the lowest binding energy (BE) in kcal/mol. BE values less than −6 kcal/mol are considered to represent significant binding affinity.

### Drug-like and Lead-like Hit Determination

Ligands that yielded a best docking pose with BE<−6 were additionally filtered using physio-chemical properties of compounds. These included lipophilicity (LogP: log of octanal-water partition coefficient) and solubility (LogS) using the SMARTS notation available from SILICOS-IT[51], which were implemented in .NET. Molecular weight (MW), topological surface area (TPSA), number of hydrogen bond donors (HBD), hydrogen bond acceptors (HBA), and number of rotatable bonds (RotB) were determined using OB’s .NET assembly. All compounds were kekulized and stripped of salts prior to calculation of physio-chemical properties, except for LogS solubility calculations, for which hydrogens were added. Two sets of criteria were employed for assessing suitability of ligands for lead discovery: “drug-like” and “lead-like”. The drug-like hits were based on the Muegge (Bayer) criteria [52] for which 200≤MW≤600, −2≤LogP≤5, TPSA≤150, HBD≤5, HBA≤10, and RotB≤15. Whereas the lead-like criteria were LogP<3, MW<300, HBD≤3, HBA≤3, and RotB≤3.

### Computational Toxicity and ADME Prediction

Determination of the absorption, distribution, metabolism, excretion (ADME) and toxicity and of new and existing drugs is necessary to identify their harmful effects of humans, animals, plants, and their environment. Historically, *in vivo* animal models have been applied for ADME and toxicity testing; however, these are constrained by time, ethical considerations, and financial burden. As an alternative, *in silico* computational methods can be used to simulate, analyze, and visualize predictions for ADME and toxicity. *In silico* ADME and toxicology predictions can complement drug design to prioritize chemicals, guide toxicity tests, and minimize late-stage failure of new drugs. Computational prediction can also potentially minimize the need for animal testing, reduce costs and time for toxicity testing, and improve toxicity and safety assessment. Early-stage identification of hazardous new compounds can also improve the cost-benefits of novel drug synthesis.

### Fathead Minnow Toxicity (FMT)

The Fathead minnow is an important aquatic and terrestrial toxicity endpoint target, and Fathead minnow toxicity data were obtained from Cheng et al. [53]. FMT toxicity data consisted of 188 FMT− and 366 FMT+ compounds (554 total). The FMT endpoint for each compound was expressed as the concentration lethal to 50% of the organisms (LC50) for FMT during 96h flow-through exposure tests. Cheng et al. [53] selected a threshold value of LC50=0.5mmol/L to partition the data into low and high acute FMT compounds. Compounds with the value of LC50 less than 0.5 mmol/L were assigned as high acute FMT compounds, whereas others were assigned as low acute FMT compounds. The chemical name, CAS numbers, FMT test results, and SMILES strings were available in the data.

### Honey Bee Toxicity (HBT)

195 pesticides or pesticide-like molecules for HBT (96 HBT−, 99 HBT+) were collected from Cheng et al. [53], based on data from the US EPA ECOTOX database[54]. The HBT endpoint for Apis mellifera bees was expressed as the dose lethal to 50% of the test population (LD50) during a 48h exposure test. Cheng et al. [53] selected a threshold value of LD50=100 μg/bee to designate high acute HBT compounds and low acute HBT compounds. Compounds with an LD50 below 100 μg/bee were coded as high acute HBT compounds, while others were coded as low acute HBT compounds. The chemical name, CAS numbers, HBT test results, and SMILES strings were available in the data.

### Tetrahymena Pyriformis Toxicity (TPT)

Tetrahymena pyriformis toxicity (TPT) is often used as a toxicology endpoint, and 1571 diverse TPT-tested chemicals were collected from Cheng et al. [55]. Toxicity data was expressed as the negative logarithm of 50% growth inhibitory concentration (pIGC50) values and duplicated molecules were removed. Xue et al. [56] selected a threshold value of pIGC50=−0.5 for discriminating TPT and non-TPT compounds (compounds with pIGC50>−0.5 were assigned as TPT, otherwise as non-TPT). The entire dataset was then divided into 1217 TPT+ and 354 TPT− compounds. The chemical name, CAS numbers, SMILES strings and pIGC50 value of 1571 compounds were available in the data.

### Human Intestinal Absorption (HIA)

The original HIA dataset was collected from Shen et al. [57]. This dataset contained n=578 compounds with fraction absorption (%FA) values. Shen et al. also specified a threshold value of 30% to partition compounds into HIA+ and HIA− (78 HIA− and 500 HIA+ compounds). Drugs with oral dosage formulations were considered to be HIA+ compounds. The chemical name, SMILES and class labels HIA+ and HIA-were available in the data.

### Blood Brain Barrier Penetration (BBB)

The BBB dataset contained n=1593 compounds, also obtained from Shen et al. [57], and have been categorized into BBB+ (n=1283) and BBB− (n=310). The chemical name, CAS numbers, BBB test results, and SMILES strings were available in the data.

### Cytochrome P450 Inhibition (CYP)

The P450 gene superfamily is involved in the metabolism of approximately 90% of approved drugs and clearance of xenobiotics. Drug safety and toxicity is directly hinged to CYP enzyme inhibition, because if a compound is a strong inhibitor of a CYP enzyme, it can result in reduced metabolism of drugs that are a substrate of the CYP enzyme, potentially leading to toxic serum plasma levels. Therefore, it is imperative to develop *in silico* prediction models of CYP inhibition for new compounds to project safety and toxicity. A large dataset containing more than 13,445 unique compounds against five major CYP isoforms, namely, 1A2, 2C9, 2C19, 2D6, and 3A4, was obtained from the PubChem AID-1851 database [58]. The assay employed for generation of these data used various human CYP P450 isozymes to measure the dealkylation of various pro-luciferin substrates to luciferin. The luciferin is then measured by luminescence after the addition of a luciferase detection reagent. Pro-luciferin substrate concentration in the assay was equal to its KM for its CYP P450 isozyme. Inhibitors and some substrates limit the production of luciferin and decrease measured luminescence. A compound was classified as a CYP inhibitor if the AC50 (the compound concentration leads to 50% of the activity of an inhibition control) value was 10 μM. A compound was considered as a non-inhibitor if AC50 was >57μM. Regarding samples sizes, for CYP1A2 there were 13,256 total compounds with 7,256 non-inhibitors and 6,000 inhibitors, for CYP2C9 there were 12,901 compounds with 8,782 non-inhibitors and 4,119 inhibitors, for CYP2C19 there were 13,445 molecules with 7,532 non-inhibitors and 5,913 inhibitors, for CYP2D6 there were 13,910 compounds 11,139 non-inhibitors and 2,771 inhibitors, and for CYP3A4 there were 13,017 compounds with 7,751 non-inhibitors and 5,266 inhibitors. The chemical name, CAS numbers, CYP test results, and SMILES strings were available in the data.

### Chemical Fingerprints for Toxicity and ADME Predictions

One approach to computational ADME and toxicity prediction employs chemical substructure analysis of known compounds which have been tested and [59] applies the associative rules between structure and outcome to new compounds whose substructure has been determined. The traditional method for identifying chemical substructure in compounds has been based on the FP2 fingerprint, which yields the presence (absence) of various atoms, bonds, aromaticity and cyclicity, and fine structure of a compound. FP2 fingerprints are in the form of binary 1024-bit vectors which signify presence and absence of the various moieties. It is important to note that while the granularity of FP2 fingerprints is high, there is less available information related to copy number of substructure elements, so any exercise is essentially hinged to a binary yes/no dilemma.

Using the toxicity and ADME training data described above, we employed the .NET OB assembly [47] to transform SMILES strings for each training compound into a FP2 1024-bit vector representing chemical substructures. OB yields FP2 fingerprints in the form of 256 4-byte Hex characters were translated to binary bits. Bit values were transformed from 0 to −1, and 1 to 1+ and appended to an analytic file with ADME or toxicity outcomes of the respective training molecule. Toxicity and ADME predictions for the selected DTP ligands were based on trained logistic regression models using 25-100 fingerprints that achieved an ROCAUC>65% for leave-one-out cross validation. Therefore, the predictive results are crude approximations.

Figure 1 illustrates the workflow employed for all ligand preparation, receptor preparation, docking, drug- and lead-like filtering of docked ligands, and computational toxicology and ADME predictions.

**Figure 1.**
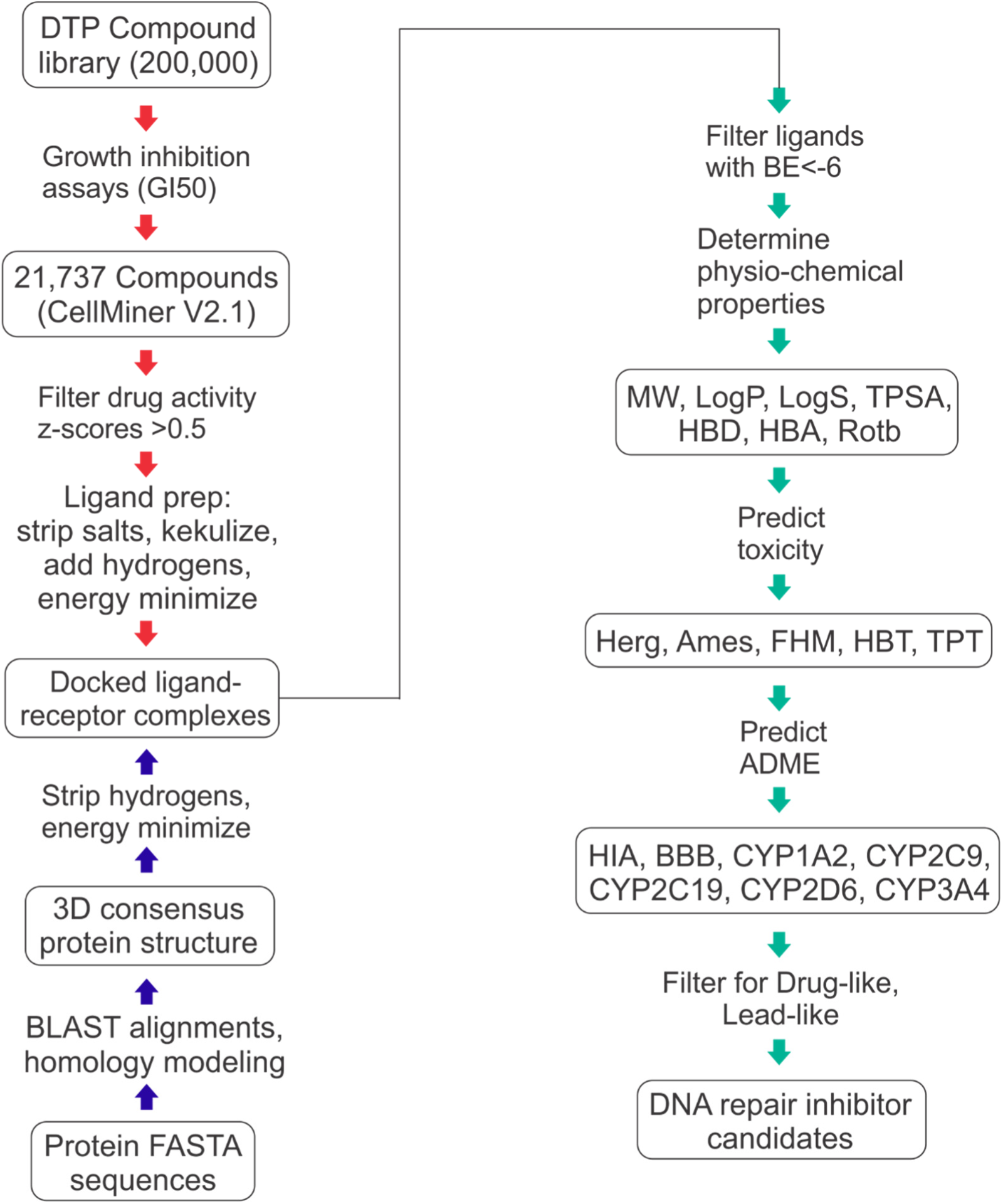
Workflow for ligand-receptor docking and drug- and lead-like hit identification.

## RESULTS

Table 2 lists the number of docked ligands as well as the number of drug-like and lead-like hits. The ligands that underwent docking were required to have a sensitive dose-response (z-score>0.5) in more than half of the DTP cell lines representing each cancer. Drug-like and lead-like criteria for physio-chemical properties were only applied to ligands whose receptor binding energies were <−6 kcal/mol for the best docking pose. For AML, among 3835 docked ligands, 1181 were drug-like and 173 were lead-like. For breast cancer, there were 684 ligands docked, for which 237 were drug-like and 47 were lead-like. GBM, on the other hand involved 343 docked ligands, for which 238 were drug-like and 38 were lead-like. Colon cancer ligand-receptor docking included 1123 ligands, with 751 being drug-like and 106 taking on lead-like properties. Lung cancer had the least number of docked ligands, with 31 identified having drug-activity, 20 that were drug-like, and 7 which were lead-like. Melanoma involved 291 docked ligands, with 224 revealing drug-like properties and 52 yielding lead-like properties. There were 105 docked ligands for ovarian cancer, with 75 drug-like and 8 lead-like. Finally, for renal papillary cancer there were 161 ligands docked, with 103 being drug-like, and 24 being lead-like.

**Table 2.** Number of docked ligands, and drug-like or lead-like ligands identified*.

| **Cancer** | **Docked** | **Drug-like** | **Lead-like** |
| --- | --- | --- | --- |
| AML | 3835 | 1181 | 173 |
| Breast | 684 | 237 | 47 |
| GBM | 343 | 238 | 38 |
| Colon | 1123 | 751 | 106 |
| Lung | 31 | 20 | 7 |
| Melanoma | 291 | 224 | 52 |
| Ovarian | 105 | 75 | 8 |
| Renal papillary | 161 | 103 | 24 |

\*Docked compounds were required to have drug activity z-scores >0.5 in more than half of the DTP cell lines tested for each cancer. Docked candidates for drug-like and lead-like filtering were required to have binding energies <-6 kcal/mol for the best docking pose.

For the cancers investigated, we were able to identify many energy-minimized compounds which were strongly bound to energy-minimized receptors. Altogether, our results indicate that many compounds were strongly bound to multiple receptors, and passed criteria for being drug-like, with fewer compounds portraying lead-like properties. The mean binding energy for ligand-receptor docking of each of the cancers considered was −7.42(1.22) for GBM, −7.19(1.2) for ovarian, −6.94(0.78) for lung, −6.93(1.25) for renal, −6.91(1.05) for melanoma, −6.88(1.04) for breast, −6.82(1.06) for colon, and −6.6(1.27) for AML. The top 10 most strongly bound receptors were PARP2(−8.88), REV1(−8.38), DDB1(−8.35), MUS81(−8.22), ALKB2(−8.01), XRCC5(−7.62), RFC3(−7.62), MUTYH(−7.5), POLI(−7.47), and FANCD2(−7.47), which revealed the importance of these receptors as potential druggable targets for therapy. Figure 2 shows the cancer-specific BE distribution for all possible ligand-receptor pairs. The large proportion of significant docking poses with BE<−6 kcal/mol are readily visible.

**Figure 2.**
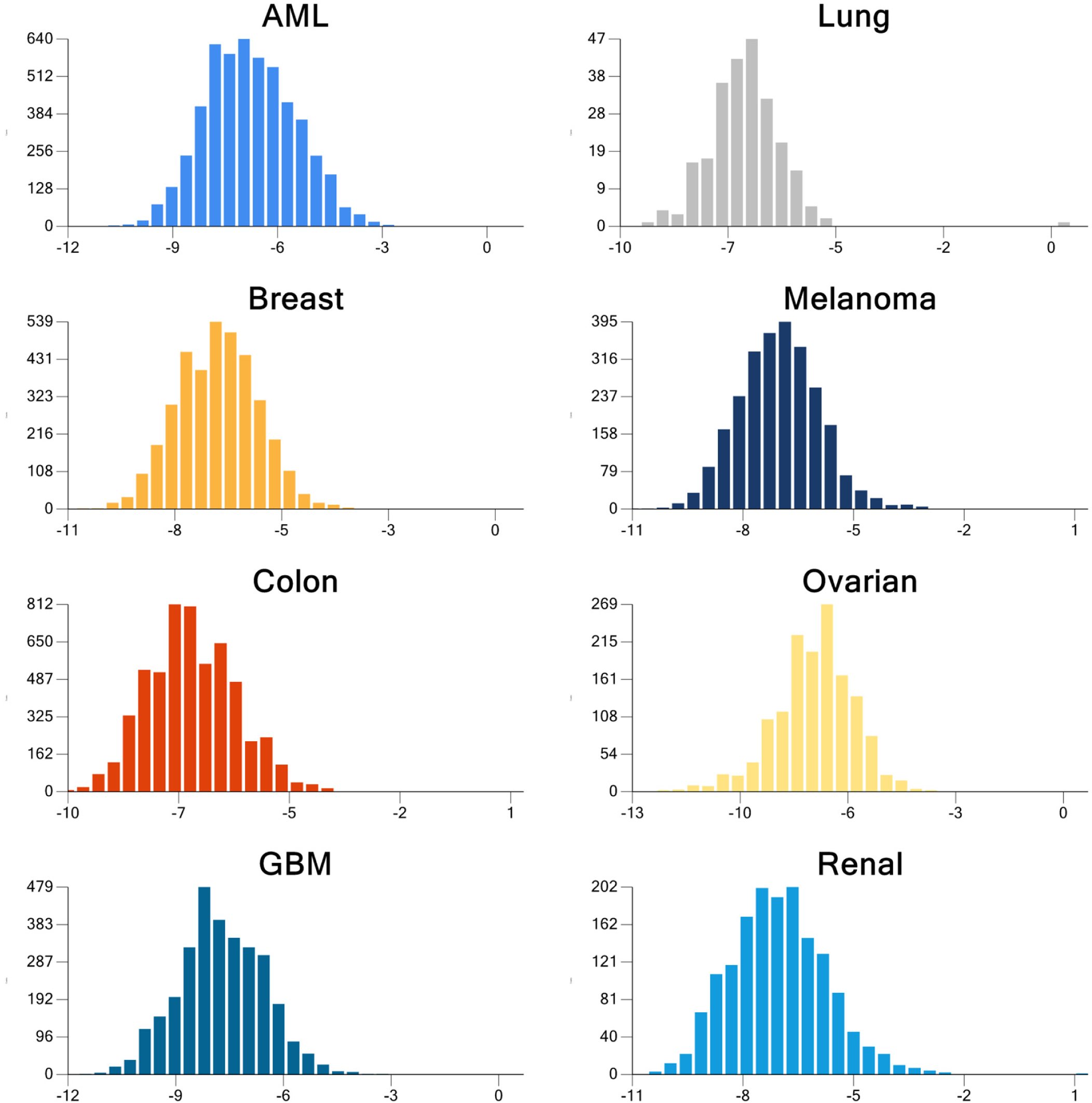
Distribution of the docking binding energy, BE (kcal/mol), from the best pose for all possible combinations of ligand-receptor pairs for each cancer. BE values <−6 kcal/mol are considered significant.

Figures 3–6 illustrate the 2D molecular structure for the top drug-like 20 ligands for AML, breast, colon, and lung cancer. Supplemental Figures S1–S4 contain 2D structure plots of the top 20 drug-like hits for GBM, melanoma, ovarian, and renal papillary cancer. Scaffold analysis for the drug-like and lead-like ligands followed by cluster analysis is now being pursued to determine if there are unique clusters of compounds. Tables 3–6 list the physio-chemical properties and computational ADME and toxicity predictions for the top 20 drug-like ligands shown in Figures 3–6. (Tables S1–S4 list these same parameters for GBM, melanoma, ovarian, and renal papillary cancers). As one can notice, lipophilicity values (LogP) fell within the acceptable range. Compounds which are too lipophilic (LogP>5) are often associated with greater metabolic clearance, metabolite-related toxicity, lower solubility, less oral absorption, and affinity for hydrophobic targets instead of the desired target, which increases promiscuity-related off-target toxicity. Low lipophilicity can also increase renal clearance, and negatively impact permeability and decrease potency, resulting in lower efficacy. Most of the ligands reported also had solubilities (LogS) near lower bound of −4 for the majority of approved drugs; compounds with LogS<−6 are classified as being poorly soluble. Poorly soluble compounds tend to have poor absorption, low stability, and fast clearance[60]. Less soluble compounds are also more difficult to handle and formulate, and parenteral delivery requires greater levels of solubility[61]. Most of the top 20 drug-like ligands appeared to be BBB permeable and readily absorbed in the intestine (i.e., HIA) as indicated by high prediction probabilities. While Herg channel blockage and the Ames carcinogenicity tests did not seem to be of too much concern, there were several ligands which resulted in high probabilities for FHM, HBT, and TPT toxicity. However, during the stages of discovery, it is customary to sacrifice false positives (lower specificity) in toxicity, while prioritizing greater sensitivity for efficacy, due to the greater uncertainty in adverse events during clinical studies. There also appeared to be wide variation in the predicted inhibition of cytochrome P-450 (CYP) enzymes, which may or may not turn out to be a metabolic or safety concern. Our future in vitro and in vivo experiments will require additional filtering within the lists of drug-like and lead-like candidates (results not shown). In addition, further toxicity and ADME predictions will be pursued to refine these estimates.

**Figure 3.**
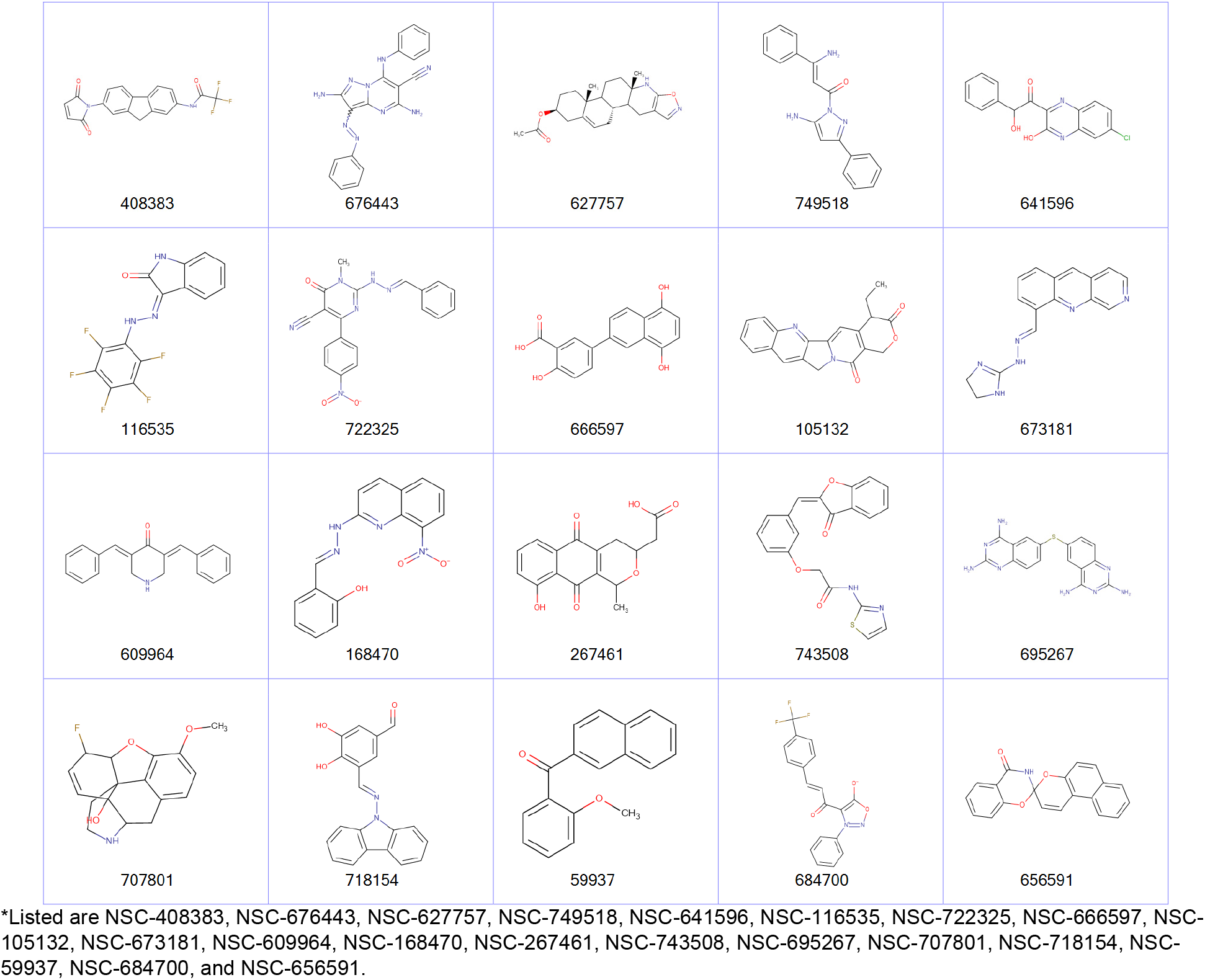
2D structures of top 20 drug-like ligands for AML*.

**Figure 4.**
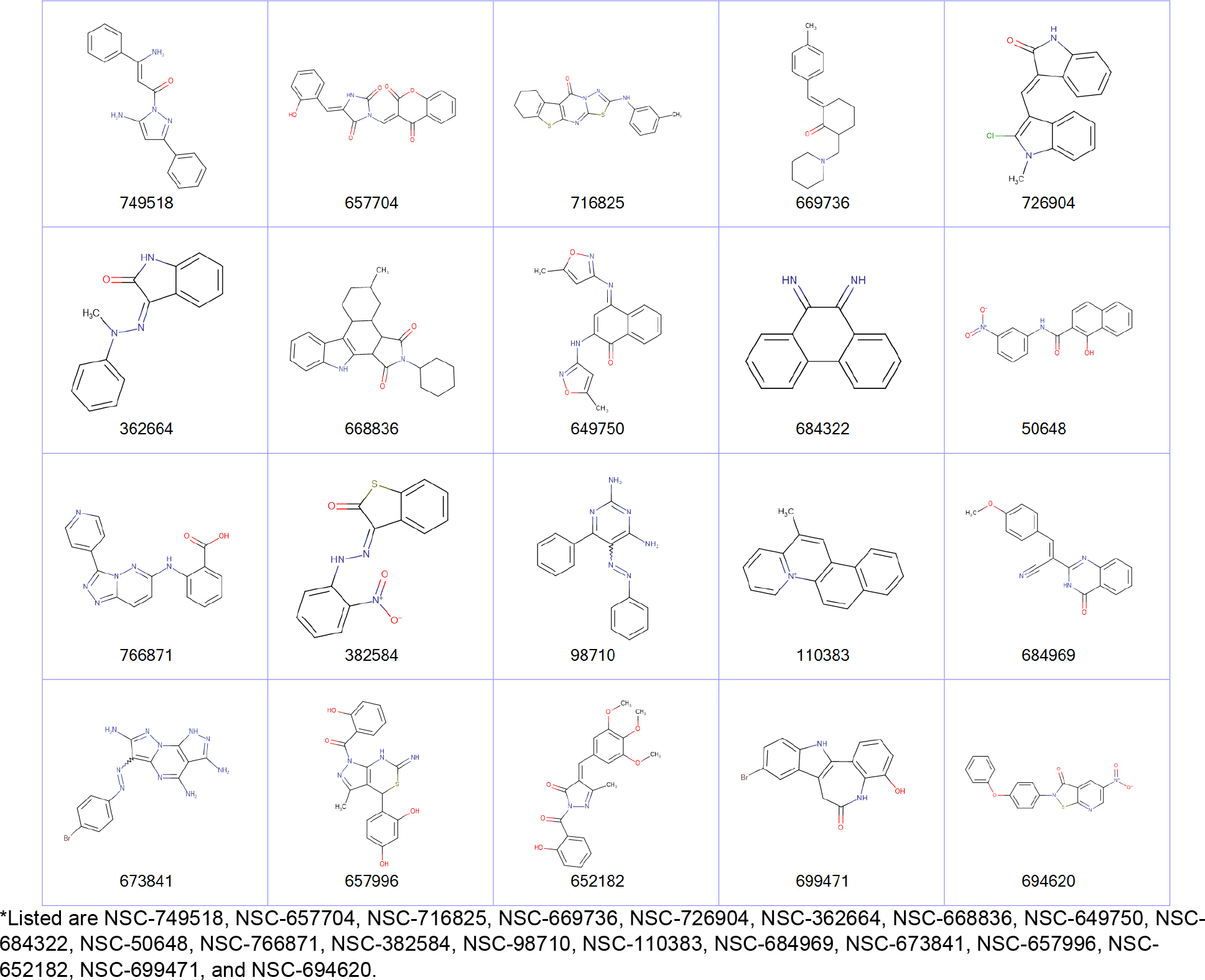
2D structures of top 20 drug-like ligands for breast cancer*.

**Figure 5.**
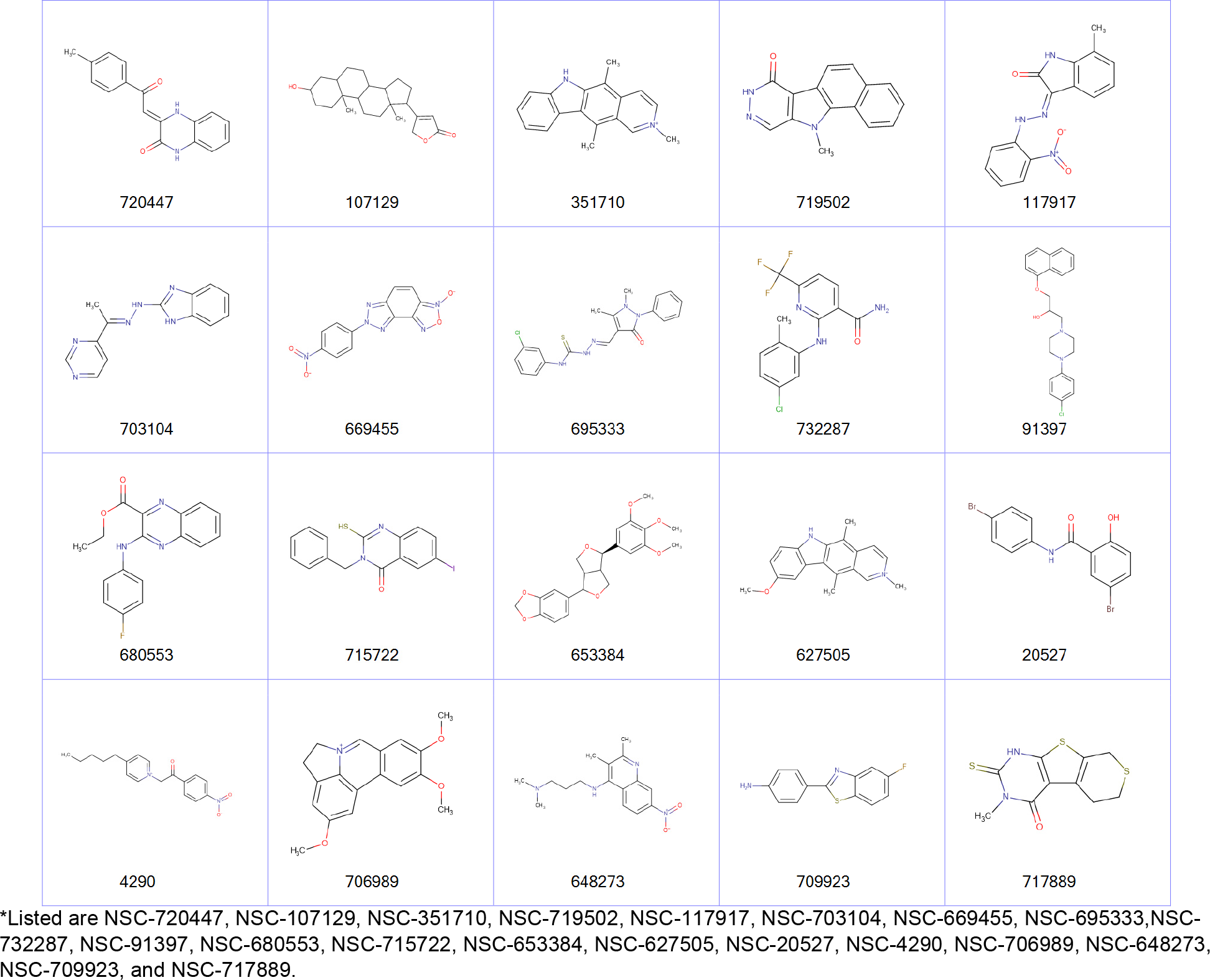
2D structures of top 20 drug-like ligands for lung cancer*.

**Figure 6.**
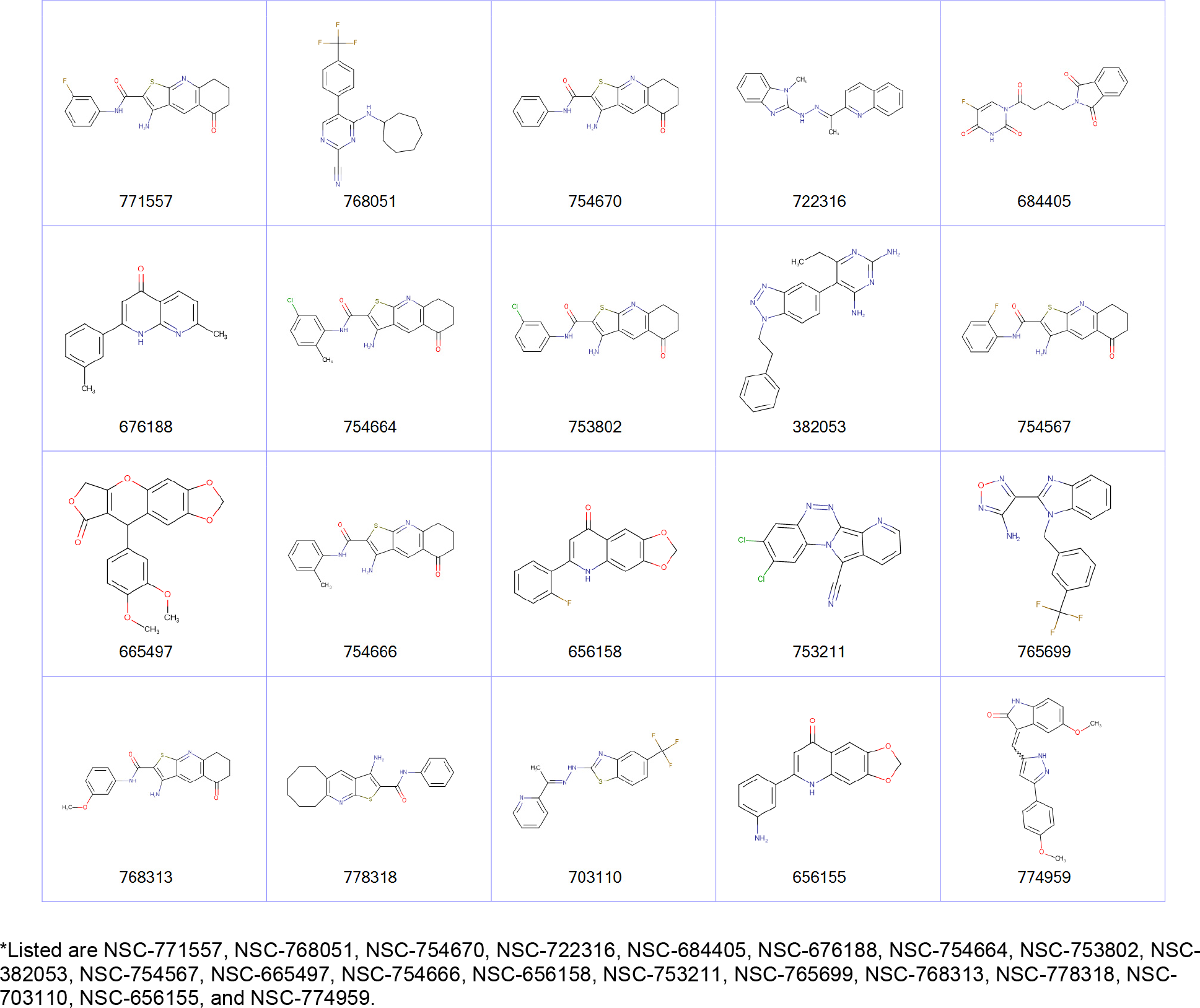
2D structures of top 20 drug-like ligands for colon cancer*.

**Table 3.** AML: List of physio-chemical properties and predicted toxicology and ADME for top 20 ligands. Toxicology and ADME predictions are probabilities in the range [0,1]*.

| Name | MW | LogP | LogS | TPSA | Rotb | HBD | HBA | BBB | HIA | HERG | AMES | FHM | HBT | TPT | CYP1A2 | CYP2C9 | CYP2C19 | CYP2D6 | CYP3A4 |
| --- | --- | --- | --- | --- | --- | --- | --- | --- | --- | --- | --- | --- | --- | --- | --- | --- | --- | --- | --- |
| 408383 | 372.3 | 3.87 | -4.67 | 66.48 | 4 | 1 | 3 | 0.99 | 0.98 | 0.16 | 0.16 | 1 | 0.24 | 0.98 | 0.68 | 0.67 | 0.66 | 0.27 | 0.42 |
| 676443 | 369.38 | -0.69 | -5.12 | 112.58 | 4 | 3 | 5 | 0.99 | 0.94 | 0.61 | 0.61 | 0.03 | 0.84 | 0 | 0.75 | 0.22 | 0.29 | 0.14 | 0.33 |
| 627757 | 370.49 | 2.29 | -3.95 | 38.33 | 2 | 1 | 2 | 1 | 0.62 | 0.01 | 0.01 | 0 | 0.8 | 0 | 0.14 | 0.13 | 0.27 | 0.16 | 0.37 |
| 749518 | 304.35 | 0.37 | -3.84 | 69.11 | 4 | 2 | 2 | 0.94 | 0.99 | 0.01 | 0.01 | 0.35 | 0.93 | 0.88 | 0.87 | 0.38 | 0.63 | 0.06 | 0.4 |
| 641596 | 314.72 | 1.33 | -4.35 | 57.53 | 3 | 2 | 5 | 1 | 1 | 0.01 | 0.01 | 1 | 0 | 0.95 | 0.4 | 0.56 | 0.6 | 0.03 | 0.24 |
| 116535 | 327.21 | 4.79 | -5.89 | 53.49 | 2 | 2 | 2 | 1 | 0.98 | 0.95 | 0.95 | 0.41 | 0.02 | 0.96 | 0.76 | 0.42 | 0.55 | 0.12 | 0.24 |
| 722325 | 374.35 | 1.96 | -5.23 | 133.11 | 5 | 1 | 4 | 0.95 | 0.98 | 0.1 | 0.1 | 0.03 | 0.14 | 0.88 | 0.85 | 0.19 | 0.4 | 0.08 | 0.31 |
| 666597 | 296.27 | 2.73 | -3.34 | 97.99 | 2 | 4 | 5 | 0.89 | 0.98 | 0.04 | 0.04 | 0.99 | 0.7 | 0.87 | 0.55 | 0.13 | 0.36 | 0.08 | 0.18 |
| 105132 | 332.35 | 3.43 | -5.2 | 43.37 | 1 | 0 | 3 | 0.93 | 0.99 | 0.35 | 0.35 | 0.36 | 0.99 | 0.95 | 0.83 | 0.57 | 0.62 | 0.28 | 0.53 |
| 673181 | 290.32 | 1.59 | -5.19 | 48.78 | 3 | 2 | 4 | 0.97 | 0.98 | 0.09 | 0.09 | 0.08 | 0.22 | 0.88 | 0.91 | 0.2 | 0.44 | 0.55 | 0.28 |
| 609964 | 275.34 | 4.47 | -5.48 | 29.1 | 2 | 1 | 2 | 1 | 0.98 | 0.13 | 0.13 | 0.79 | 0.93 | 0.98 | 0.49 | 0.22 | 0.42 | 0.36 | 0.25 |
| 168470 | 308.29 | 2.46 | -4.86 | 112.48 | 4 | 2 | 3 | 0.73 | 0.98 | 0.59 | 0.59 | 0.34 | 0.29 | 0.89 | 0.96 | 0.35 | 0.49 | 0.09 | 0.32 |
| 267461 | 302.28 | 2.43 | -2.04 | 100.9 | 2 | 2 | 6 | 0.69 | 0.96 | 0.01 | 0.01 | 1 | 0.98 | 0.94 | 0.18 | 0.3 | 0.4 | 0.04 | 0.41 |
| 743508 | 378.4 | 1.54 | -5.51 | 64.63 | 6 | 1 | 5 | 0.81 | 0.9 | 0.01 | 0.01 | 0.92 | 0.81 | 0.95 | 0.93 | 0.79 | 0.75 | 0.07 | 0.8 |
| 695267 | 350.4 | -2.92 | -4.37 | 129.38 | 2 | 4 | 4 | 1 | 0.95 | 0.36 | 0.36 | 0.98 | 0.49 | 0.93 | 0.76 | 0.14 | 0.5 | 0.3 | 0.46 |
| 707801 | 303.33 | 3.3 | -2.68 | 50.72 | 1 | 2 | 4 | 1 | 0.82 | 0.55 | 0.55 | 0.77 | 0.97 | 0.94 | 0.05 | 0.43 | 0.2 | 0.53 | 0.75 |
| 718154 | 330.34 | 3.74 | -4.53 | 69.89 | 3 | 2 | 4 | 0.96 | 0.96 | 0.07 | 0.07 | 0.9 | 0.37 | 0.88 | 0.86 | 0.3 | 0.45 | 0.1 | 0.24 |
| 59937 | 262.3 | 4.73 | -5.89 | 26.3 | 3 | 0 | 2 | 1 | 0.98 | 0.09 | 0.09 | 0.8 | 0.7 | 0.98 | 0.77 | 0.18 | 0.37 | 0.26 | 0.23 |
| 684700 | 360.29 | 0.13 | -4.94 | 40.13 | 5 | 0 | 3 | 0.95 | 0.99 | 0.57 | 0.57 | 0.73 | 0.15 | 0.96 | 0.62 | 0.24 | 0.65 | 0.09 | 0.75 |
| 656591 | 315.32 | 4.39 | -5.67 | 47.56 | 0 | 1 | 3 | 0.83 | 1 | 0.09 | 0.09 | 0.01 | 0.77 | 0.87 | 0.61 | 0.34 | 0.78 | 0.23 | 0.61 |

\*Listed are NSC-408383, NSC-676443, NSC-627757, NSC-749518, NSC-641596, NSC-116535, NSC-722325, NSC-666597, NSC-105132, NSC-673181, NSC-609964, NSC-168470, NSC-267461, NSC-743508, NSC-695267, NSC-707801, NSC-718154, NSC-59937, NSC-684700, and NSC-656591. Receptor binding energies, promiscuity, genotoxicity, skin sensitivity, and aquatic toxicity from SMARTS hits not listed.

**Table 4.**
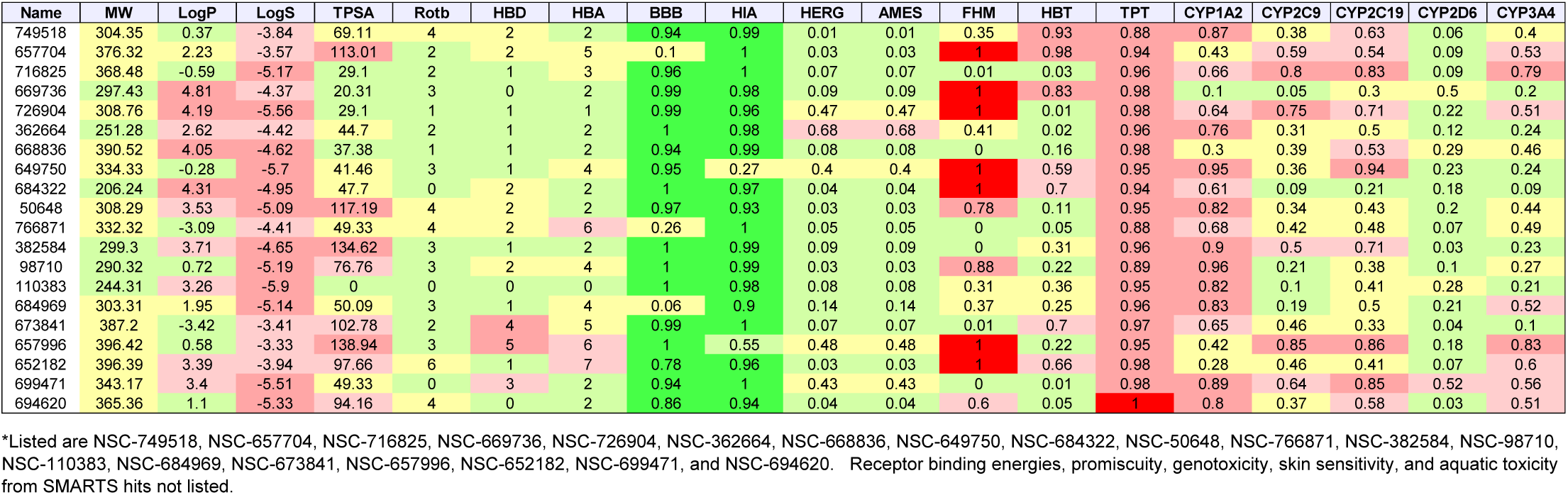
Breast cancer: List of physio-chemical properties and predicted toxicology and ADME for top 20 ligands. Toxicology and ADME predictions are probabilities in the range [0,1]*.

| Name | MW | LogP | LogS | TPSA | Rotb | HBD | HBA | BBB | HIA | HERG | AMES | FHM | HBT | TPT | CYP1A2 | CYP2C9 | CYP2C19 | CYP2D6 | CYP3A4 |
| --- | --- | --- | --- | --- | --- | --- | --- | --- | --- | --- | --- | --- | --- | --- | --- | --- | --- | --- | --- |
| 749518 | 304.35 | 0.37 | -3.84 | 69.11 | 4 | 2 | 2 | 0.94 | 0.99 | 0.01 | 0.01 | 0.35 | 0.93 | 0.88 | 0.87 | 0.38 | 0.63 | 0.06 | 0.4 |
| 657704 | 376.32 | 2.23 | -3.57 | 113.01 | 2 | 2 | 5 | 0.1 | 1 | 0.03 | 0.03 | 1 | 0.98 | 0.94 | 0.43 | 0.59 | 0.54 | 0.09 | 0.53 |
| 716825 | 368.48 | -0.59 | -5.17 | 29.1 | 2 | 1 | 3 | 0.96 | 1 | 0.07 | 0.07 | 0.01 | 0.03 | 0.96 | 0.66 | 0.8 | 0.83 | 0.09 | 0.79 |
| 669736 | 297.43 | 4.81 | -4.37 | 20.31 | 3 | 0 | 2 | 0.99 | 0.98 | 0.09 | 0.09 | 1 | 0.83 | 0.98 | 0.1 | 0.05 | 0.3 | 0.5 | 0.2 |
| 726904 | 308.76 | 4.19 | -5.56 | 29.1 | 1 | 1 | 1 | 0.99 | 0.96 | 0.47 | 0.47 | 1 | 0.01 | 0.98 | 0.64 | 0.75 | 0.71 | 0.22 | 0.51 |
| 362664 | 251.28 | 2.62 | -4.42 | 44.7 | 2 | 1 | 2 | 1 | 0.98 | 0.68 | 0.68 | 0.41 | 0.02 | 0.96 | 0.76 | 0.31 | 0.5 | 0.12 | 0.24 |
| 668836 | 390.52 | 4.05 | -4.62 | 37.38 | 1 | 1 | 2 | 0.94 | 0.99 | 0.08 | 0.08 | 0 | 0.16 | 0.98 | 0.3 | 0.39 | 0.53 | 0.29 | 0.46 |
| 649750 | 334.33 | -0.28 | -5.7 | 41.46 | 3 | 1 | 4 | 0.95 | 0.27 | 0.4 | 0.4 | 1 | 0.59 | 0.95 | 0.95 | 0.36 | 0.94 | 0.23 | 0.24 |
| 684322 | 206.24 | 4.31 | -4.95 | 47.7 | 0 | 2 | 2 | 1 | 0.97 | 0.04 | 0.04 | 1 | 0.7 | 0.94 | 0.61 | 0.09 | 0.21 | 0.18 | 0.09 |
| 50648 | 308.29 | 3.53 | -5.09 | 117.19 | 4 | 2 | 2 | 0.97 | 0.93 | 0.03 | 0.03 | 0.78 | 0.11 | 0.95 | 0.82 | 0.34 | 0.43 | 0.2 | 0.44 |
| 766871 | 332.32 | -3.09 | -4.41 | 49.33 | 4 | 2 | 6 | 0.26 | 1 | 0.05 | 0.05 | 0 | 0.05 | 0.88 | 0.68 | 0.42 | 0.48 | 0.07 | 0.49 |
| 382584 | 299.3 | 3.71 | -4.65 | 134.62 | 3 | 1 | 2 | 1 | 0.99 | 0.09 | 0.09 | 0 | 0.31 | 0.96 | 0.9 | 0.5 | 0.71 | 0.03 | 0.23 |
| 98710 | 290.32 | 0.72 | -5.19 | 76.76 | 3 | 2 | 4 | 1 | 0.99 | 0.03 | 0.03 | 0.88 | 0.22 | 0.89 | 0.96 | 0.21 | 0.38 | 0.1 | 0.27 |
| 110383 | 244.31 | 3.26 | -5.9 | 0 | 0 | 0 | 0 | 1 | 0.98 | 0.08 | 0.08 | 0.31 | 0.36 | 0.95 | 0.82 | 0.1 | 0.41 | 0.28 | 0.21 |
| 684969 | 303.31 | 1.95 | -5.14 | 50.09 | 3 | 1 | 4 | 0.06 | 0.9 | 0.14 | 0.14 | 0.37 | 0.25 | 0.96 | 0.83 | 0.19 | 0.5 | 0.21 | 0.52 |
| 673841 | 387.2 | -3.42 | -3.41 | 102.78 | 2 | 4 | 5 | 0.99 | 1 | 0.07 | 0.07 | 0.01 | 0.7 | 0.97 | 0.65 | 0.46 | 0.33 | 0.04 | 0.1 |
| 657996 | 396.42 | 0.58 | -3.33 | 138.94 | 3 | 5 | 6 | 1 | 0.55 | 0.48 | 0.48 | 1 | 0.22 | 0.95 | 0.42 | 0.85 | 0.86 | 0.18 | 0.83 |
| 652182 | 396.39 | 3.39 | -3.94 | 97.66 | 6 | 1 | 7 | 0.78 | 0.96 | 0.03 | 0.03 | 1 | 0.66 | 0.98 | 0.28 | 0.46 | 0.41 | 0.07 | 0.6 |
| 699471 | 343.17 | 3.4 | -5.51 | 49.33 | 0 | 3 | 2 | 0.94 | 1 | 0.43 | 0.43 | 0 | 0.01 | 0.98 | 0.89 | 0.64 | 0.85 | 0.52 | 0.56 |
| 694620 | 365.36 | 1.1 | -5.33 | 94.16 | 4 | 0 | 2 | 0.86 | 0.94 | 0.04 | 0.04 | 0.6 | 0.05 | 1 | 0.8 | 0.37 | 0.58 | 0.03 | 0.51 |

\*Listed are NSC-749518, NSC-657704, NSC-716825, NSC-669736, NSC-726904, NSC-362664, NSC-668836, NSC-649750, NSC-684322, NSC-50648, NSC-766871, NSC-382584, NSC-98710, NSC-110383, NSC-684969, NSC-673841, NSC-657996, NSC-652182, NSC-699471, and NSC-694620. Receptor binding energies, promiscuity, genotoxicity, skin sensitivity, and aquatic toxicity from SMARTS hits not listed.

**Table 5.** Lung cancer: List of physio-chemical properties and predicted toxicology and ADME for top 20 ligands. Toxicology and ADME predictions are probabilities in the range [0,1]*.

| Name | MW | LogP | LogS | TPSA | Rotb | HBD | HBA | BBB | HIA | HERG | AMES | FHM | HBT | TPT | CYP1A2 | CYP2C9 | CYP2C19 | CYP2D6 | CYP3A4 |
| --- | --- | --- | --- | --- | --- | --- | --- | --- | --- | --- | --- | --- | --- | --- | --- | --- | --- | --- | --- |
| 720447 | 278.31 | 2.32 | -5.3 | 34.14 | 2 | 2 | 2 | 0.99 | 0.94 | 0.01 | 0.01 | 1 | 0.99 | 0.98 | 0.85 | 0.53 | 0.36 | 0.02 | 0.25 |
| 107129 | 358.51 | 4.7 | -2.77 | 46.53 | 1 | 1 | 2 | 0.99 | 0.94 | 0.02 | 0.02 | 1 | 0.97 | 0.69 | 0.05 | 0.02 | 0.12 | 0.02 | 0.04 |
| 351710 | 261.34 | 2.56 | -5.83 | 0 | 0 | 1 | 0 | 1 | 0.98 | 0.6 | 0.6 | 0.18 | 0.27 | 0.95 | 0.77 | 0.16 | 0.45 | 0.48 | 0.21 |
| 719502 | 249.27 | 1.22 | -4.63 | 17.07 | 0 | 1 | 2 | 0.83 | 0.98 | 0.28 | 0.28 | 0.96 | 0.27 | 1 | 0.61 | 0.18 | 0.2 | 0.08 | 0.18 |
| 117917 | 296.28 | 3.33 | -4.99 | 121.35 | 3 | 2 | 2 | 1 | 0.98 | 0.51 | 0.51 | 0.76 | 0.02 | 0.95 | 0.88 | 0.34 | 0.46 | 0.04 | 0.34 |
| 703104 | 252.27 | -1.7 | -4.47 | 24.39 | 3 | 2 | 4 | 1 | 0.69 | 0.04 | 0.04 | 0.99 | 0.21 | 0 | 0.85 | 0.16 | 0.45 | 0.35 | 0.14 |
| 669455 | 298.21 | -5.19 | -2.71 | 90.92 | 2 | 0 | 4 | 0.99 | 0.38 | 0.04 | 0.04 | 1 | 0.35 | 0.92 | 0.65 | 0.15 | 0.27 | 0.05 | 0.21 |
| 695333 | 399.9 | 3.17 | -5.01 | 85.58 | 6 | 2 | 2 | 0.16 | 0.78 | 0.3 | 0.3 | 0 | 0.05 | 0 | 0.41 | 0.67 | 0.84 | 0.05 | 0.7 |
| 732287 | 329.7 | 2.53 | -5.06 | 55.12 | 4 | 2 | 2 | 1 | 0.98 | 0.14 | 0.14 | 0 | 0.71 | 0.88 | 0.82 | 0.45 | 0.68 | 0.08 | 0.7 |
| 91397 | 396.91 | 4.31 | -5.77 | 35.94 | 6 | 1 | 3 | 0.87 | 0.95 | 0.17 | 0.17 | 0.34 | 0.38 | 0.97 | 0.45 | 0.34 | 0.53 | 0.62 | 0.33 |
| 680553 | 311.31 | 1.64 | -5.71 | 38.33 | 5 | 1 | 3 | 0.98 | 1 | 0.01 | 0.01 | 0.16 | 0.02 | 0 | 0.94 | 0.34 | 0.39 | 0.02 | 0.3 |
| 715722 | 394.23 | 3.01 | -5.16 | 55.87 | 2 | 0 | 2 | 0.64 | 0.89 | 0.49 | 0.49 | 0.88 | 0.27 | 0.95 | 0.79 | 0.14 | 0.3 | 0.3 | 0.59 |
| 653384 | 400.42 | 4.02 | -4.01 | 64.61 | 5 | 0 | 7 | 1 | 0.99 | 0.07 | 0.07 | 1 | 0.94 | 0.94 | 0.46 | 0.33 | 0.39 | 0.17 | 0.33 |
| 627505 | 291.37 | 2.58 | -5.85 | 9.23 | 1 | 1 | 1 | 0.95 | 0.98 | 0.58 | 0.58 | 0.4 | 0.27 | 0.95 | 0.75 | 0.25 | 0.58 | 0.47 | 0.31 |
| 20527 | 371.02 | 3.44 | -4.89 | 49.33 | 3 | 2 | 2 | 0.97 | 0.98 | 0.03 | 0.03 | 0.48 | 0.11 | 0.95 | 0.63 | 0.39 | 0.49 | 0.22 | 0.35 |
| 4290 | 313.37 | 2.88 | -5.13 | 84.93 | 8 | 0 | 1 | 1 | 0.98 | 0.38 | 0.38 | 0.18 | 0.27 | 0.98 | 0.64 | 0.37 | 0.45 | 0.19 | 0.33 |
| 706989 | 296.34 | 2.68 | -4.59 | 27.69 | 3 | 0 | 3 | 0.97 | 0.96 | 0.24 | 0.24 | 0.21 | 0.27 | 0.98 | 0.73 | 0.34 | 0.6 | 0.54 | 0.56 |
| 648273 | 302.37 | 2.77 | -4.94 | 83.13 | 6 | 1 | 2 | 0.99 | 0.98 | 0.35 | 0.35 | 0 | 0.16 | 0.98 | 0.83 | 0.17 | 0.48 | 0.71 | 0.65 |
| 709923 | 244.29 | 1.46 | -4.69 | 26.02 | 1 | 1 | 1 | 1 | 0.97 | 0.06 | 0.06 | 0.18 | 0.37 | 0.89 | 0.85 | 0.34 | 0.69 | 0.1 | 0.38 |
| 717889 | 270.39 | 1.82 | -2.31 | 74.46 | 0 | 1 | 1 | 0.99 | 0.94 | 0.04 | 0.04 | 0 | 0.4 | 0.5 | 0.51 | 0.45 | 0.52 | 0.08 | 0.11 |

\*Listed are NSC-720447, NSC-107129, NSC-351710, NSC-719502, NSC-117917, NSC-703104, NSC-669455, NSC-695333, NSC-732287, NSC-91397, NSC-680553, NSC-715722, NSC-653384, NSC-627505, NSC-20527, NSC-4290, NSC-706989, NSC-648273, NSC-709923, and NSC-717889. Receptor binding energies, promiscuity, genotoxicity, skin sensitivity, and aquatic toxicity from SMARTS hits not listed.

**Table 6.** Colon cancer: List of physio-chemical properties and predicted toxicology and ADME for top 20 ligands. Toxicology and ADME predictions are probabilities in the range [0,1]*.

| Name | MW | LogP | LogS | TPSA | Rotb | HBD | HBA | BBB | HIA | HERG | AMES | FHM | HBT | TPT | CYP1A2 | CYP2C9 | CYP2C19 | CYP2D6 | CYP3A4 |
| --- | --- | --- | --- | --- | --- | --- | --- | --- | --- | --- | --- | --- | --- | --- | --- | --- | --- | --- | --- |
| 771557 | 355.39 | 1.83 | -5.31 | 72.19 | 3 | 2 | 3 | 1 | 0.98 | 0.43 | 0.43 | 0 | 0.02 | 0.98 | 0.79 | 0.72 | 0.77 | 0.37 | 0.93 |
| 768051 | 360.38 | 2.59 | -5.79 | 35.82 | 4 | 1 | 3 | 1 | 0.99 | 0.07 | 0.07 | 0.47 | 0.45 | 0.96 | 0.89 | 0.12 | 0.57 | 0.45 | 0.76 |
| 754670 | 337.4 | 1.41 | -5.09 | 72.19 | 3 | 2 | 3 | 1 | 0.98 | 0.26 | 0.26 | 0 | 0.02 | 0.98 | 0.81 | 0.62 | 0.74 | 0.38 | 0.93 |
| 722316 | 315.37 | 1.37 | -5.86 | 24.39 | 3 | 1 | 3 | 0.99 | 0.88 | 0.49 | 0.49 | 0 | 0.2 | 0.95 | 0.82 | 0.27 | 0.37 | 0.39 | 0.22 |
| 684405 | 345.28 | 0.75 | -3.2 | 88.59 | 5 | 1 | 5 | 0.42 | 0.98 | 0.04 | 0.04 | 0 | 0.59 | 0.95 | 0.47 | 0.32 | 0.4 | 0.08 | 0.33 |
| 676188 | 250.3 | 2.64 | -5.79 | 17.07 | 1 | 1 | 2 | 0.9 | 0.98 | 0.29 | 0.29 | 0.02 | 0.06 | 0.89 | 0.9 | 0.13 | 0.46 | 0.2 | 0.4 |
| 754664 | 385.87 | 2.56 | -5.92 | 72.19 | 3 | 2 | 3 | 1 | 0.98 | 0.27 | 0.27 | 0 | 0 | 0.98 | 0.81 | 0.81 | 0.87 | 0.38 | 0.94 |
| 753802 | 371.84 | 2.05 | -5.59 | 72.19 | 3 | 2 | 3 | 1 | 0.98 | 0.36 | 0.36 | 0 | 0 | 0.98 | 0.81 | 0.81 | 0.87 | 0.38 | 0.93 |
| 382053 | 359.43 | -1.54 | -5.77 | 52.04 | 5 | 2 | 4 | 0.88 | 0.98 | 0.07 | 0.07 | 1 | 0.01 | 0.95 | 0.69 | 0.48 | 0.56 | 0.31 | 0.49 |
| 754567 | 355.39 | 1.83 | -5.31 | 72.19 | 3 | 2 | 3 | 1 | 0.98 | 0.55 | 0.55 | 0 | 0.02 | 0.98 | 0.79 | 0.62 | 0.74 | 0.4 | 0.93 |
| 665497 | 368.34 | 4.23 | -4.27 | 72.45 | 3 | 0 | 6 | 0.93 | 0.99 | 0.18 | 0.18 | 1 | 0.98 | 0.94 | 0.34 | 0.45 | 0.68 | 0.33 | 0.41 |
| 754666 | 351.42 | 1.92 | -5.43 | 72.19 | 3 | 2 | 3 | 1 | 0.98 | 0.19 | 0.19 | 0 | 0.02 | 0.98 | 0.81 | 0.63 | 0.72 | 0.38 | 0.94 |
| 656158 | 283.25 | 3.7 | -5.31 | 35.53 | 1 | 1 | 3 | 0.96 | 0.97 | 0.46 | 0.46 | 0.68 | 0.18 | 0.95 | 0.92 | 0.23 | 0.74 | 0.39 | 0.69 |
| 753211 | 314.13 | 0.53 | -5.05 | 23.79 | 0 | 0 | 4 | 0.96 | 1 | 0.02 | 0.02 | 0.78 | 0.66 | 0.86 | 0.73 | 0.35 | 0.46 | 0.16 | 0.3 |
| 765699 | 359.31 | -1.11 | -5.41 | 26.02 | 4 | 1 | 3 | 0.46 | 0.63 | 0.26 | 0.26 | 0.02 | 0.16 | 0.88 | 0.86 | 0.39 | 0.73 | 0.46 | 0.63 |
| 768313 | 367.42 | 1.46 | -5.11 | 81.42 | 4 | 2 | 4 | 1 | 0.98 | 0.39 | 0.39 | 0 | 0.02 | 0.98 | 0.81 | 0.72 | 0.72 | 0.42 | 0.94 |
| 778318 | 351.47 | 2.19 | -5.71 | 55.12 | 3 | 2 | 2 | 1 | 0.98 | 0.62 | 0.62 | 0 | 0.02 | 0.98 | 0.71 | 0.64 | 0.73 | 0.4 | 0.91 |
| 703110 | 336.33 | 1.03 | -5.5 | 24.39 | 4 | 1 | 3 | 1 | 0.74 | 0.11 | 0.11 | 0 | 0.08 | 0.96 | 0.86 | 0.33 | 0.67 | 0.08 | 0.32 |
| 656155 | 280.28 | 2.56 | -4.68 | 61.55 | 1 | 2 | 3 | 0.81 | 0.97 | 0.42 | 0.42 | 0.68 | 0.22 | 0.95 | 0.94 | 0.23 | 0.76 | 0.44 | 0.73 |
| 774959 | 347.37 | 2.1 | -5.97 | 47.56 | 4 | 2 | 4 | 0.81 | 0.99 | 0.26 | 0.26 | 0.04 | 0.18 | 0.98 | 0.72 | 0.71 | 0.74 | 0.13 | 0.66 |

\*Listed are NSC-771557, NSC-768051, NSC-754670, NSC-722316, NSC-684405, NSC-676188, NSC-754664, NSC-753802, NSC-382053, NSC-754567, NSC-665497, NSC-754666, NSC-656158, NSC-753211, NSC-765699, NSC-768313, NSC-778318, NSC-703110, NSC-656155, and NSC-774959. Receptor binding energies, promiscuity, genotoxicity, skin sensitivity, and aquatic toxicity from SMARTS hits not listed.

## DISCUSSION

The adjustment of gene expression for the PCNA metagene in oncology is not a new concept[32]; however, there are groups which are unaware of its existence, and the novel perspective that can be attained once it is performed. Approximately 50% of the genome is co-regulated by PCNA, and because its effect is so widespread, it is imperative that investigations in cancer research account for and remove the effects of this non-cancer related transcriptional mechanism. Our effort to remove PCNA effects from RNA-Seq based gene expression in TCGA data is novel, and this has resulted in new lists of DNA repair genes which are predictive of OS beyond chance variation. Regarding molecular docking, virtual drug screens typically employ hundreds of thousands of ligands for which no dose-response information is available prior to the analysis. We took the opposite approach by filtering on cell line-derived drug activity z-scores, and only used DTP compounds which elicited a sensitive dose-response. Therefore, the only remaining uncertainties surrounding the strongly binding ligands are whether the observed DTP cell line sensitivities were due to a direct/indirect inhibitory mechanism, underlying toxicity independent of binding, or both. These issues can be addressed in future in vitro experiments to confirm protein binding and in vivo xenograft models to confirm animal model efficacy.

The results support repair inhibition models of cancer survival, which can be applied to longitudinal studies and clinical trials. The molecular docking employed revealed results that are potentially hinged to repair inhibition via protein binding. Overall, our docking analysis allowed us to make inferences about cell line sensitivity to draw conclusions about the relationships between binding and tumor growth for common cancers. The translational value of our results is established by the identification of drug-like and lead-like compounds. This could prove useful in future studies of molecular markers of therapy for delayed progression. Our future investigations will start with the top 50–100 ligand identified for each cancer to link verified gene-knockdown with xenograft models for confirming an association between repair inhibition and experimental data for tumor growth and survival. Molecular docking of DTP compounds with DNA repair proteins has enabled us to gain a perspective that hit identification and repair inhibition in the cell line data employed could likely reveal new compounds and mechanisms for oncotherapy. This view will hopefully enforce an appreciation among oncologists and biologists for the translational value of DTP cell line testing results, compounds, and molecular docking of these compounds with DNA repair proteins, for potential clinical trials involving single-or multi-label treatments associated with prolonged survival and pursuing longitudinal studies to improve therapeutic strategies.

We did not comparatively assess gene expression in the tumor cell lines employed. Recall that although the DTP project has used a variety of gene expression platforms, expression patterns in cell lines was not available for most of the DNA repair genes used in this investigation. We also did not employ a gold-standard to establish false positive and false negative rates of toxicological and ADME endpoints. Rather, we ranked ligands by their binding energies hinged to MD and approximated computationally the toxicological and ADME outcomes. Experimental validation of the binding efficiency of ligands and in vitro determination of delayed tumor growth, toxicological, and ADME outcomes are needed in order to assess whether select groups of patients will benefit from new therapies arising from our findings. The work presented here suggests that molecular docking studies of DNA repair proteins with cell line-sensitive DTP compounds can provide new insights into the development of human cancer and can establish new leads for future research on molecular diagnosis and therapeutics.

There are several challenging issues surrounding development of tumor progression models based on repair inhibition. First, there is the problem of inherited germline polymorphisms, which confer a variety of risks and require a variety of treatment regimens. Second, tumor heterogeneity and immune escape are others hallmarks of cancer that cannot be easily overcome when searching for new modes of therapy. The RNA-Seq based data obtained from TCGA are not based on single-cell analysis, which would be helpful for elucidating heterogeneity; however, the large genetic variation identified throughout the TCGA samples used would exacerbate the complexity surrounding our attempt to portray repair inhibition via a single picture. Third, our approach was bioinformatic and not mechanistic using laboratory methods, since the intent was purely computational at this stage of investigation. We also did not consider DNA methylation status, chromosome aberrations, and microsatellite instability, which would overlay more complexity on the results obtained.

In conclusion, our bioinformatic approach to infer repair inhibition from cell line and RNA-Seq data in the absence of data on genomic alterations and germline polymorphisms should be cautiously interpreted. While most human cancers are nevertheless sporadic, the importance of the inherited component of cancer, combined with genomic alterations and tumor heterogeneity leading to growth and progression was not addressed in this investigation.

## CONCLUSION

We employed computational methods to derive ligand-receptor binding and prediction of toxicological and ADME endpoints for synthetic and natural compounds in the DTP repository and DNA repair proteins which were predictive of OS after adjustment for the PCNA metagene. Results of our computational methods translate to portraits of potentially new repair inhibitors of delayed tumor progression. The utility of our findings can be realized by oncologists and biologists who envision new targets and mechanisms of repair inhibition. We conclude that the results presented can be directly employed for in vitro and in vivo confirmatory experiments to identify new targets of therapy for cancer survival.

## ABBREVIATIONS

ADME: Absorption, distribution, metabolism, and excretion
AML: Acute myelogenous leukemia
BBB: Blood brain barrier
BE: Binding energy (kcal/mol)
CNS: Central nervous system
CYP: Cytochrome P-450 enzymes
GBS: Glioblastoma multiforme
HERG: Human ether-a-go-go related gene
HBA: Hydrogen bond acceptors
HBD: Hydrogen bond donors
HIA: Human intestinal absorption
LogS: Solubility
LogP: Lipophilicity
MW: Molecular weight (Daltons)
MD: Molecular dynamics
PCNA: Proliferating cell nuclear antigen
RotB: Number of rotatable bonds (covalent)
TCGA: The Cancer Genome Atlas
TPSA: Topological surface area

## ETHICS APPROVAL AND CONSENT TO PARTICIPATE

The list of DNA repair genes that were used were identified in a separate external investigation; however, the TCGA data are publicly available at cBio-Portal (http://www.cbioportal.org)

## AVAILABILITY OF DATA

The TCGA human genomic data that are publicly available at cBio-Portal (http://www.cbioportal.org).

A list of all human DNA repair genes can be found at https://www.mdanderson.org/documents/Labs/Wood-Laboratory/human-dna-repair-genes.html

The DTP compounds are available at https://discover.nci.nih.gov/cellminer/

## COMPETING INTERESTS

The author declares no competing interests.

## AUTHOR CONTRIBUTION

LEP wrote the manuscript, developed the workflow, performed the homology modeling, ligand and receptor preparation, docking, toxicity and ADME prediction.

## FUNDING

Research was supported by NASA Grant NNX-12AO52A.

